# Nanopore sequencing reveals *U2AF1 S34F*-associated full-length isoforms

**DOI:** 10.1101/871863

**Authors:** Cameron M. Soulette, Eva Hrabeta-Robinson, Alison Tang, Maximillian G. Marin, Angela N. Brooks

## Abstract

*U2AF1 S34F* is one of the most recurrent splicing factor mutations in lung adenocarcinoma (ADC) and has been shown to cause transcriptome-wide pre-mRNA splicing alterations. While *U2AF1 S34F*-associated splicing alterations have been described, the function of altered mRNA isoform changes remains largely unexplored. To better understand the impact *U2AF1 S34F* has on isoform fate and function, we conducted high-throughput long-read cDNA sequencing from isogenic human bronchial epithelial cells with and without *U2AF1 S34F* mutation. We found that nearly 75% (49,366) of our long-read constructed multiexon isoforms do not overlap GENCODE or short-read assembled isoforms. We found 198 transcript isoforms with significant expression and usage changes caused by *U2AF1 S34F* mutation, including a novel lncRNA. Isoforms from immune-related genes were largely downregulated in mutant cells, none of which were found to have splicing changes. Finally, isoforms likely targeted by nonsense-mediated decay were preferentially downregulated in *U2AF1 S34F* cells, suggesting that the impact of observed isoform changes may alter the translational output of affected genes. Altogether, long-read sequencing provided additional insights into transcriptome alterations and downstream functional consequences associated with *U2AF1 S34F* mutation.

## Introduction

Previous cancer genomic studies across lung adenocarcinoma (ADC) patients have revealed recurrent mutations in the splicing factor *U2AF1 (Brooks et al., 2014; Cancer Genome Atlas Research Network, 2014; Campbell et al., 2016)*. U2AF1 is an essential splicing factor that functions to identify the 3’ end of intronic sequence in the early steps of pre-mRNA splicing (Shao *et al.*, 2014). In ADC, the most recurrent *U2AF1* mutation occurs at amino acid residue 34, in which a C>T transition causes a change from serine to phenylalanine (S34F). The impact of *U2AF1 S34F* on pre-mRNA splicing has been widely studied (Przychodzen *et al.*, 2013; Brooks *et al.*, 2014; Coulon *et al.*, 2014; Ilagan *et al.*, 2015; Shirai *et al.*, 2015; Park *et al.*, 2016; Yip *et al.*, 2017; Palangat *et al.*, 2019; Smith *et al*., 2019), and previous work has shown that mutant U2AF1 has an altered binding affinity with its pre-mRNA substrate (Okeyo-Owuor *et al.*, 2015; Fei *et al.*, 2016). In ADC, mutant *U2AF1* has been shown to alter pre-mRNA splicing and other post-transcriptional processes (Brooks *et al.*, 2014; Fei *et al.*, 2016; Park *et al.*, 2016; Palangat *et al.*, 2019).

The impact of *U2AF1* mutations on the transcriptome raises interesting hypotheses for an oncogenic role through mRNA dysregulation. *U2AF1 S34F* is known to alter alternative-splicing and polyadenylation of cancer-relevant genes (Przychodzen *et al.*, 2013; Brooks *et al.*, 2014; Ilagan *et al.*, 2015; Okeyo-Owuor *et al.*, 2015; Shirai *et al.*, 2015; Yip *et al.*, 2015, 2017; Fei *et al.*, 2016; Park *et al.*, 2016; Smith *et al.*, 2019). In myelodysplastic syndromes, *U2AF1 S34F* alters pre-mRNA splicing of Interleukin-1 receptor-associated kinase 4 (IRAK4), producing isoforms that promote activation of kappa-light-chain-enhancer of B cells (NF-kB), a factor known to promote leukemic cell growth (Smith *et al.*, 2019). In addition to splicing-dependent functions of *U2AF1 S34F*, recent studies show a splicing-independent post-transcriptional role for *U2AF1 S34F* in modulating translational efficiency of genes involved in inflammation and metastasis in human bronchial epithelial cells (Palangat *et al*., 2019). Although some oncogenic roles for *U2AF1 S34F* have been described, the full functional impact of *U2AF1*-associated mRNAs are unknown.

Investigating mRNA isoform function proves difficult given the complexity and accuracy of isoform assembly with short-reads (Engström *et al.*, 2013; Steijger *et al.*, 2013). Accurate isoform assembly is important in investigating RNA processing alterations associated with global splicing factors, like U2AF1. Recent studies have shown the utility of long-read approaches in capturing full-length mRNA isoforms, by constructing isoforms missed by short-read assembly methods (Oikonomopoulos *et al.*, 2016; Byrne *et al.*, 2017; de Jong *et al.*, 2017; Tang *et al.*, 2018; Workman *et al.*, 2019). Moreover, long-read approaches have already been conducted using RNA derived from primary tumor samples harboring *SF3B1* mutations, demonstrating its effectiveness in capturing mutant splicing factor transcriptome alterations *(Tang et al., 2018)*. In addition, studies have shown the extent to which long-read data can be used as a quantitative measure for gene expression (Oikonomopoulos *et al.*, 2016; Byrne *et al.*, 2017). Given the global impact of *U2AF1* mutations on the transcriptome, identifying RNA processing alterations at the level of full-length mRNA isoforms is an essential step in understanding the functional impact of affected mRNAs.

Here we used an emerging long-read sequencing approach to characterize isoform structure and function of transcript isoforms affected by *U2AF1 S34F*. We chose to study *U2AF1 S34F*-associated isoform changes in an isogenic cell line, HBEC3kt cells, which has been used as a model for identifying transcriptome changes associated with *U2AF1 S34F* (Ramirez et. al. 2004, Fei et. al. 2016). We constructed a long-read transcriptome that contains substantial novel mRNA isoforms not reflected in annotations, nor could they be reconstructed using short-read sequencing assembly approaches. Our long-read data supports a strong *U2AF1 S34F* splicing phenotype, in which we demonstrate the ability to recapitulate the splicing phenotype associated with *U2AF1 S34F* mutants using splicing event-level analyses. We find that isoforms containing premature termination codons (PTCs) and immune-related genes are significantly downregulated. Finally, we leverage previously published short-read polysome profiling data to show changes in translational control for genes affected by *U2AF1 S34F*. Our work provides additional insights into the function of transcripts altered by *U2AF1 S34F* mutation.

## Results

### Long-read sequencing reveals the complexity of the HBEC3kt transcriptome

We first characterized the transcriptome complexity of HBEC3kt cells with and without *U2AF1 S34F* mutation using the Oxford Nanopore minION platform. We conducted cDNA sequencing on two clonal cell lines, two wild-type and two *U2AF1 S34F* mutation isolates (WT1, WT2, MT1, MT2). We extracted whole-cell RNA from each cell isolate, one growth replicate of WT1 and M1 and two replicates of WT2 and MT2. We converted RNA into cDNA using methods described in previous nanopore sequencing studies ((Picelli *et al.*, 2013; Byrne *et al.*, 2017)**; Methods**), and performed nanopore 1D cDNA sequencing on individual flow cells (**Figure 1A**). Our sequencing yielded 8.8 million long-reads across all 6 sequencing runs (**Supplemental Table 1**), which we subsequently processed through a Full-Length Alternative Isoform analysis of RNA (FLAIR; (Tang *et al.*, 2018) to construct a reference transcriptome and perform various differential analyses (**Figure 1B, Methods**). We constructed a total of 63,289 isoforms, 49,366 of which were multi-exon and 45,749 contained unique junction sets (**Supplemental Figure 1, Supplemental File 1**).

We compared FLAIR isoforms against GENCODE reference annotations (v19) and a short-read assembly using previously published data from HBEC3kt cells (Fei *et al.*, 2016)**; Supplemental File 2; Methods**). We found that only one-third of our FLAIR transcriptome overlapped with GENCODE annotations (**Figure 2A)**. The remaining FLAIR isoforms contained novel elements, such as novel exons, novel junction combination, or a novel genomic locus. In contrast, nearly half of the isoforms from short-read assembly were comprised of known GENCODE isoforms. We hypothesized that the increased number of annotated isoforms from short-read assembly could be due to higher sequencing depths. We therefore overlapped intron junction-chains between all three datasets and quantified expression from each overlapping group (**Figure 2B**). We found significant differences in expression for isoforms not contained in our set of high confident FLAIR isoforms (p-value <0.001; **Figure 2B** top panel).

**Figure 1.**
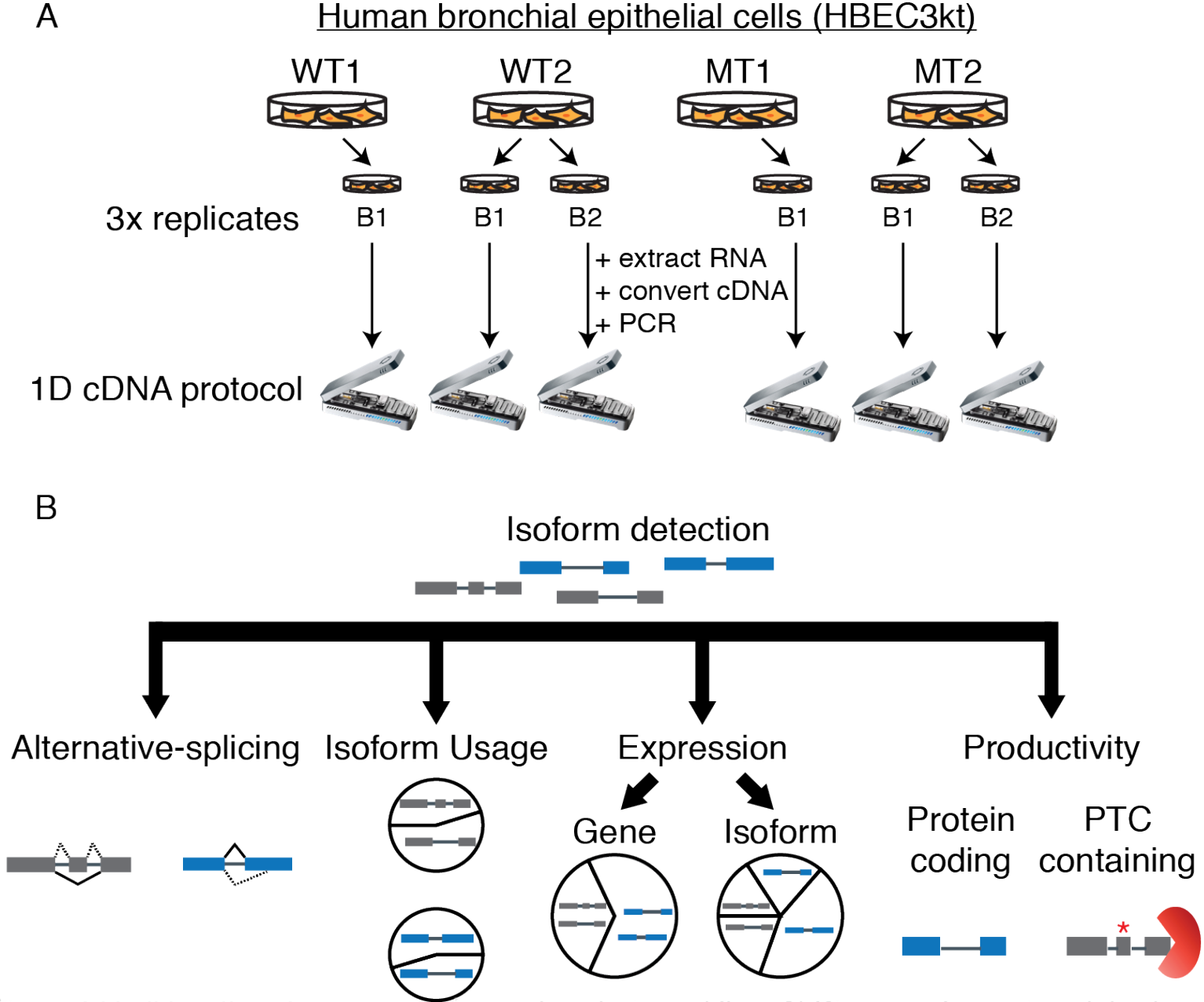
Full-length isoform sequencing and analysis workflow. A) Diagram of experimental setup and sequencing strategy. RNA was extracted from whole cell lysate and converted to cDNA using a polyA tail selection strategy. Wild-type and mutant conditions were sequenced in triplicate. Each sequencing run was conducted in parallel, in which a wild-type or mutant was sequenced on separate flow cells. B) Data processing pipeline workflow. FLAIR was used to construct a reference transcriptome from long-read data with matched short-read RNA-seq and to perform differential expression and productivity analyses.

Although our long-read approach did not capture lowly expressed isoforms, we found a large proportion of FLAIR-exclusive isoforms that contained novel exons, junction combinations and novel loci isoforms (**Figure 2C**). Notably, we identified 182 FLAIR-exclusive isoforms from 123 unannotacted loci, none of which were assembled by short-reads despite having short-read coverage support, perhaps due to repeat elements that are known to be difficult to assemble across (Treangen & Salzberg 2012). We investigated a putative lncRNA we call *USFM* (<u>u</u>pregulated in <u>s</u>plicing <u>f</u>actor <u>m</u>utant; *LINC02879*), which was one of the most highly expressed multi-exon isoforms in mutant samples (**Figure 2C,** bottom panel). We manually examined long-reads aligned to *USFM* and found poly(A) tails, suggesting *USFM* supporting reads are not likely to be 3’ end fragmented products. Next, we used publicly available ENCODE data to look for transcription factor binding sites and histone modification marks (ENCODE Project Consortium, 2012) that would provide additional evidence of a transcribed gene locus. Peaks associated with H3K27 acetylation and H3K4 methylation suggest the presence of regulated transcribed genomic regions. Moreover, the transcript start site of *USFM* overlapped with active promoter predictions from chromHMM (ENCODE Project Consortium, 2012), an algorithm used to predict promoters and transcriptionally active regions (**Figure 2C,** bottom panel). No significant homology matches to protein-coding domains could be found using NCBI BLAST (Johnson *et al.*, 2008). Taken together, these data indicate that *USFM* isoforms have characteristics that are consistent with lncRNAs, and highlight the utility of long-reads in identifying putative novel genes.

**Figure 2.**
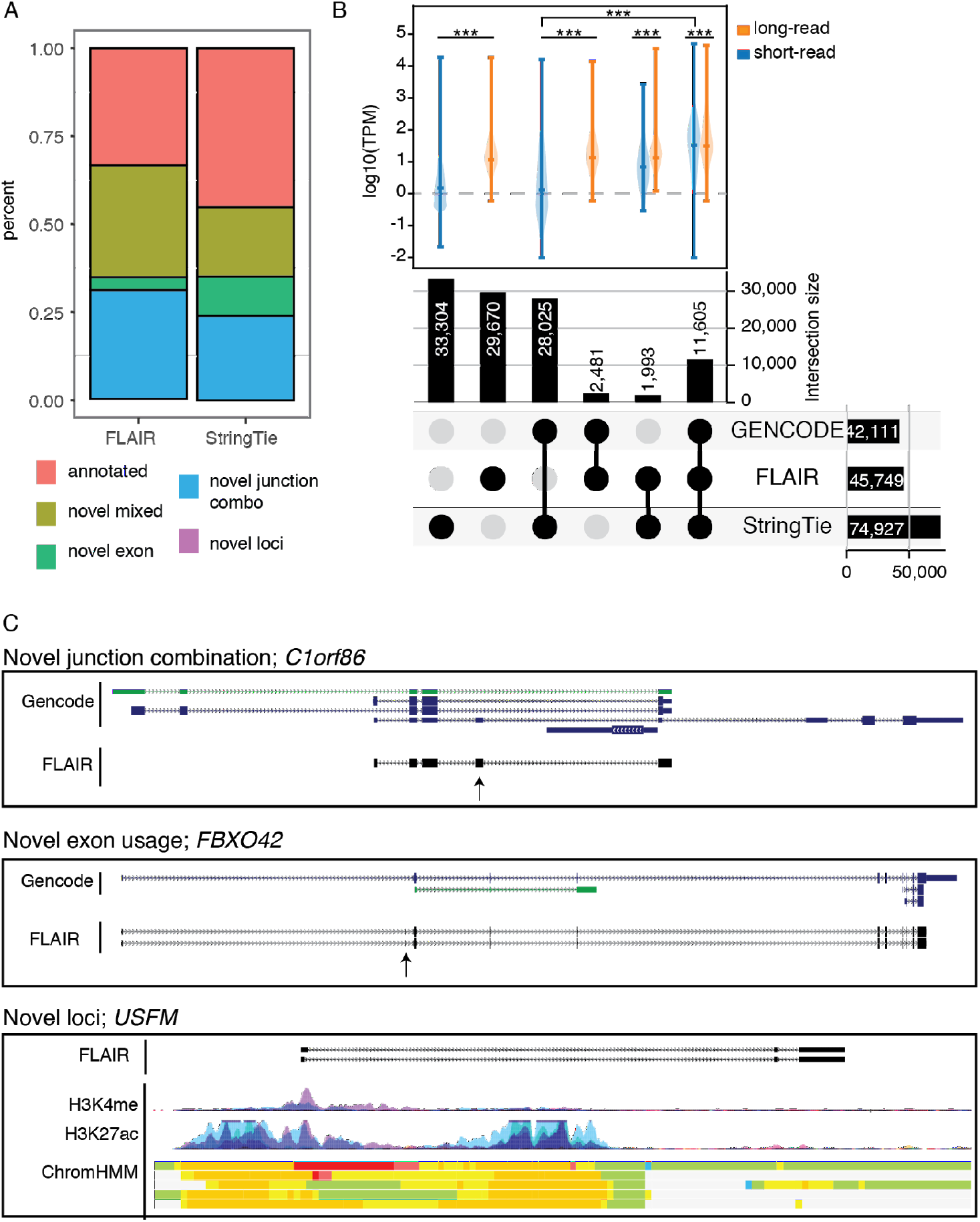
FLAIR captures HBEC3kt transcriptome complexity. A)Isoform annotation categories for FLAIR and StringTie isoforms in comparison to GENCODE v19 annotations. B) Overlap of transcript isoforms between long-read FLAIR, short-read StringTie assembly, and Gencode v19 annotation. Top panel, expression quantification from StringTie and FLAIR-quantify for isoforms in each overlap category. Expression distributions were compared using Wilcoxon rank sum test, and comparisons denoted with *** have p-values < 0.001. C) UCSC genome browser shot example for novel classification categories. For each panel, Gencode annotations represent gencode v19 basic annotation set. For Novel Exon and Novel Loci panels, Encode regulatory tracks were included to show H3K27 acetylation, DNase hypersensitivity Transcription factor binding ChIP (TF Chip), and ChromHMM data from various cell lines. Red and yellow hues represent putative promoter regions; Green regions represent putative transcribed regions.

### *U2AF1 S34F* splicing signature captured by long-read event-level analyses

We compared *U2AF1 S34F*-associated splicing signatures in our long-read data to those found from analyses of short-read datasets (Brooks *et al.*, 2014; Ilagan *et al.*, 2015; Okeyo-Owuor *et al.*, 2015; Fei *et al.*, 2016). Previous reports have shown that cassette exon skipping is the most prevalent splicing alteration induced by *U2AF1 S34F*. Moreover, motif analysis of 3’ splice sites adjacent to altered cassette exons with enhanced and reduced inclusion show a strong nucleotide context for ‘CAG’ and ‘TAG’ acceptor sites, respectively (Brooks *et al.*, 2014; Ilagan *et al.*, 2015; Okeyo-Owuor *et al.*, 2015; Fei *et al.*, 2016). As shown previously, we find that cassette exon events are the most predominant patterns of altered splicing associated with *U2AF1 S34F* mutations in short-read sequencing from The Cancer Genome Atlas (TCGA) lung ADC samples and HBEC3kt isogenic cell lines (179/226 and 187/225, respectively; (Fei *et al.*, 2016). We observe minor differences in alternative donor and intron retention events between TCGA and HBEC3kt short-read, which could likely be explained by the limitations of statistical testing with only two replicates for the HBEC3kt short-read data (Hansen *et al.*, 2011).

We next assayed for alternative-splicing alterations in our long-read data using FLAIR-diffSplice (**Figure 3A**). We used FLAIR for our event-level long-read analysis since most alternative-splicing algorithms were designed for short-read analysis (**Supplemental Table 4**; **Methods**). Our long-read event-level analysis was consistent with short-read analysis. (**Figure 3A**). The most predominant altered splicing pattern from long-read data was cassette exons, in which we found a general trend toward exon exclusion (51/55; **Supplemental Figure 2A**). In addition, we found a good correlation in the magnitude of change in percent spliced in (PSI) between short and long-read PSI values (**Figure 3B;** Pearson r=0.88). Last, we investigated the 3’ splice site motif associated with altered cassette exons and alternative acceptor events and found ‘TAG’ and ‘CAG’ motifs associated with acceptor sites with reduced and enhanced inclusion, respectively (**Figure 3C**). Overall, our results demonstrate consistent splicing signature patterns between short and long-read methodologies.

*U2AF1 S34F* has been implicated in widespread altered poly(A) site selection (Park et. al. 2016). We took advantage of the reduced ambiguity long-reads provide in identifying poly(A) cleavage sites and identified alternative poly(A) alterations associated with *U2AF1 S34F* (**Figure 3D**). Identifying poly(A) sites with short-reads is computationally difficult due to alignment of reads primarily composed of poly(A) sequence, or alignment across repetitive sequence commonly found in 3’ untranslated regions (Chen, Ara and Gautheret, 2009; Elkon, Ugalde and Agami, 2013; Shenker *et al.*, 2015; Ha, Blencowe and Morris, 2018). We first investigated the presence of poly(A) cleavage site motifs at the 3’ ends of FLAIR isoforms, and found a strong signal ~20 nucleotides upstream from transcript end sites for the most commonly used cleavage motif, “AATAAA”, relative to random 6-mer (**Supplemental Figure 2B**). We next tested for APA site usage alterations by comparing the proportion of polyA site usage for each gene between *U2AF1* wild-type and S34F (**Methods**). 10 genes demonstrated significant changes in polyadenylation site usage (corrected p-value <0.05 & ΔAPA > 10%), which comprises 7.2% of all RNA processing alterations identified in this study(11 APA & 142 alternative-splicing events), far less than previous reports. The most significant APA alteration occurred in *BUB3* (**Figure 3E**), which is part of the mitotic checkpoint pathway, a pathway containing genes that are commonly altered in select lung cancers (Takahashi *et al*., 1999; Haruki *et al.*, 2001). Collectively, our event-level analyses confirmed our ability to capture well-documented *U2AF1 S34F*-associated splicing signatures with long-read data.

**Figure 3.**
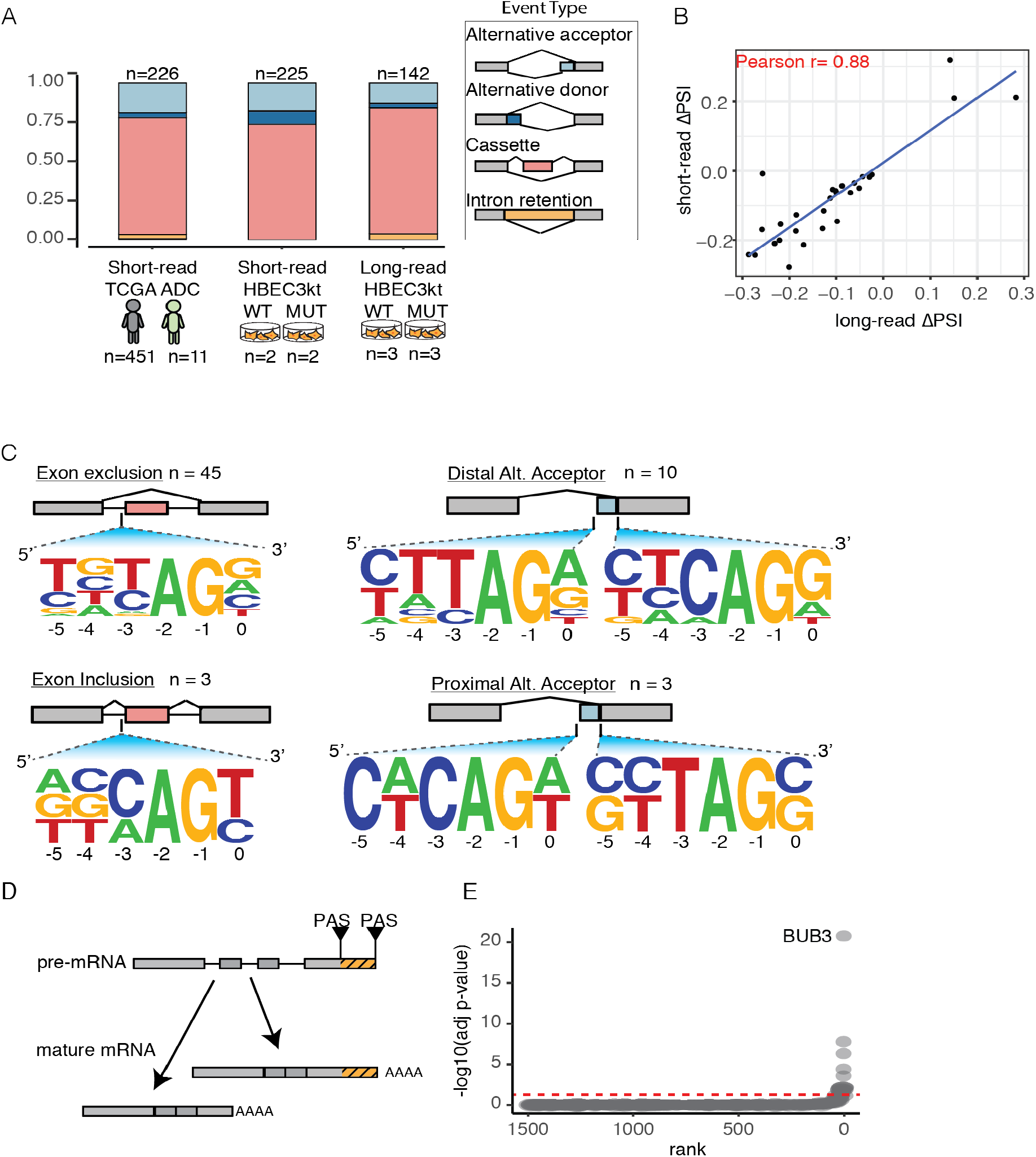
Nanopore data recapitulates *U2AF1* S34Fsplicing signature. A) Alternative-splicing events that were found to be significantly altered between wild-type and U2AF1 S34F conditions. Events are broken down into different patterns of alternative-splicing B) Change in PSI (percent spliced-in) correlation between short and long-read cassette exon events C) Weblogos of 3’ splice sites for altered cassette exons (left panels) and alternative acceptor sites (right panels) identified using nanopore data. D) Alternative polyadenylation (APA) site selection schematic. E) Ranked genes with significant changes in APA site usage.

### Long-reads provide isoform context for *UPP1* splicing alterations missed by short-read assembly

We compared the exon connectivity of cassette exons altered by *U2AF1 S34F* in Uridine phosphorylase 1 (*UPP1*), which was the most significantly altered gene in our event-level analysis. *UPP1* altered cassette exons accounted for 4 of the 55 significantly altered cassette exons (exons 5, 6-long, 6-short, and 7), one of which, exon 7, was also found to be significantly altered in TCGA ADC data (**Supplemental Table 3**). We compared 28 FLAIR isoforms containing exon 7 against StringTie assembly to determine which isoforms were missed by either method. Despite minor differences in transcript start and end sites, we found all 4 short-read assembled *UPP1* isoforms containing exon 7 in our set of FLAIR isoforms (**Supplemental Figure 2C**). The additional 21 FLAIR-exclusive isoforms contained a mixture of exon skipping events, alternative 3’ and alternative 5’ splicing events that coincided with exon 7 inclusion. A more broad comparison of all 95 *UPP1* FLAIR isoforms revealed that only 7 were assembled by short-read. We then asked if any of the FLAIR-exclusive *UPP1* isoforms were expressed at substantial proportions (>5%) by quantifying the expression of each isoform using our long-reads. We found that 6 of the 7 most highly expressed *UPP1* isoforms were FLAIR-exclusive (**Supplemental Figure 2D**). Taken together, although short-read methods assembled complex splicing regulation observed in *UPP1*, our long-read analysis revealed extensive isoform diversity not captured by short-reads.

### U2AF1 *S34F* induces strong isoform switching in *UPP1* and *BUB3*

We next assessed transcriptome-wide changes in isoform usage using our long-read data. Short-read event-level analyses typically represent isoforms by distinct RNA processing events such as splicing (**Figure 4A**). In contrast, long-reads capture entire mRNA isoforms and therefore can be used to more accurately quantify distinct isoforms. We identified 166 isoforms with significant changes in usage (corrected p-value <0.05) using DRIMSeq (**Supplemental Table 5**, **Methods**). We found that nearly half of significantly altered isoforms (82/166) had large changes in magnitude (Δisoform usage > 10%) (**Figure 4B**). Consistent with our event-level analysis, we found isoforms from *BUB3* and *UPP1* in the top 10 most significantly altered genes, suggesting that changes in these isoforms are defined by splicing event changes. Gene set enrichment analysis using the molecular signature database (Liberzon *et al.*, 2011) on differentially used isoforms (FDR<0.05, Δisoform usage > 10%) revealed genes involved in RNA metabolism, and RNA processing (**Supplemental Table 5**).

We found complex 3’ end processing patterns that define *BUB3* isoforms. Previous reports have described alternative acceptor site usage for *BUB3* that leads to the usage of distinct polyadenylation sites (Bava *et al*., 2013). Consistent with previous reports, we find that the proximal acceptor site leads to the production of isoforms using three APA sites (APA1, 2 and 3), and usage of a distal acceptor site leading to the usage of two APA sites (APA 4, 5) (**Supplemental Figure 3A**). Our event-level analyses revealed a significant shift toward usage of the distal acceptor site and APA site (Δalt. acceptor >30%; corrected p-value < 0.05), which is consistent with differences in isoform usage (**Supplemental Figure 3B**). Notably, the proximal *BUB3* 3’ acceptor site is preceded by a thymidine residue, which could partially explain isoform shifting toward the usage of the distal acceptor site. We also observed a preference for APA site 2 in wild-type samples (ΔAPA usage 20%), which seems to be lost in mutant samples (ΔAPA usage 6%; two-sided t-test p-value <0.05).

**Figure 4.**
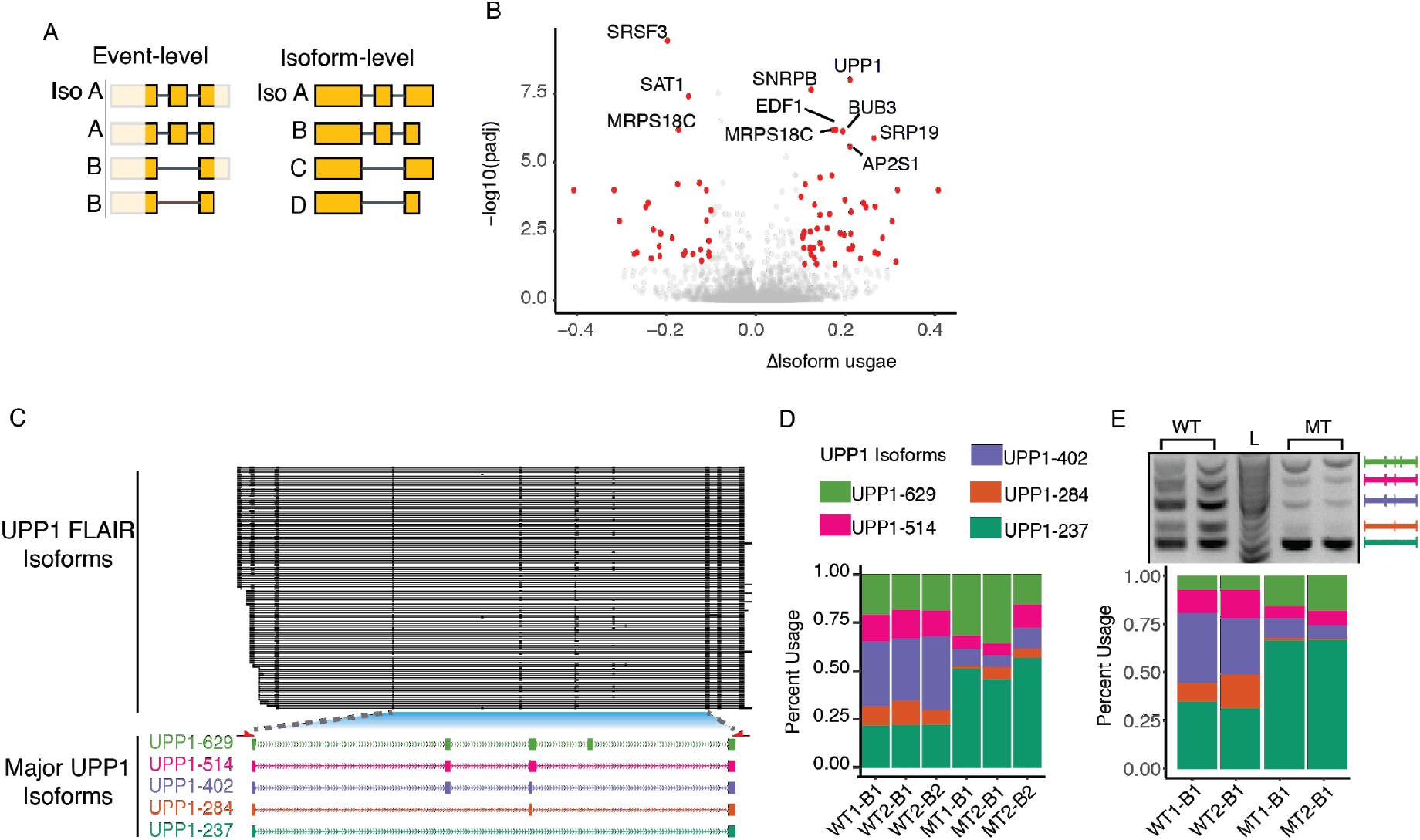
*U2AF1* S34F-associated full-length isoform usage changes. A) Diagram of event-level versus isoform-level analyses captured by long-read sequencing. alterations. B) Volcano plot of differentially used isoforms. Red dots indicate usage changes with corrected p value<0.05 and magnitude change >10%. Gene names indicate top 10 genes with significant isoform changes. C) UPP1 FLAIR isoforms (top panel), and major isoforms (bottom panel). Isoform numbers correspond to predicted amplion sizes. Red arrows below major isoforms represent PCR primers used for RT-PCR validation. D) Long-read isoform usage quantified by nanopore data. E) 2% agarose gel with UPP1 amplicon products (top panel) and gel quantification bar chart (bottom panel).

We next investigated significant isoform usage changes in *UPP1* (**Figure 4C**). Out of the 95 *UPP1* isoforms identified by our data, 68 (71%) fell below 1% of the total *UPP1* gene abundance, indicating that the majority of isoforms are minor isoforms. The remaining 28 *UPP1* isoforms were tested for differential isoform usage, 2 of which were found to have significant usage changes (corrected p-value <0.05 & Δisoform usage >10%) (**Figure 4D showing top 5 expressed isoforms**). RT-PCR validation using primers that span all *U2AF1 S34F*-associated cassette exons showed a pattern consistent with sequencing results, in which *U2AF1 S34F* induces a shift toward *UPP1* isoforms that either contain or exclude all cassette exons (**Figure 4E**). *UPP1* is known to be highly expressed in solid tumors (Lie et. al. 1998; Kanzaki et. al. 2002), but cancer associated splicing alterations have not been described. Altogether, results from our full-length isoform analysis are consistent with results from our event-level analysis, but differentiate which isoforms are actually affected by *U2AF1 S34F*.

### Isoforms changes are partially explained by event-level splicing changes

We next determined the extent to which *U2AF1 S34F* alters the expression of individual isoforms. This analysis complements our isoform switching analysis by allowing for the identification of minor isoforms (isoform usage <10%) with large expression changes, and genes with uniform isoform expression changes. Our analysis yielded 122 isoforms with significant changes in expression (corrected p-value<0.05 and log2FoldChange>1.5; **Figure 5A, Supplemental Table 6**). We found the most upregulated isoforms were from the putative lncRNA *USFM* (log2FoldChange > 3 & corrected p-value <0.01). We searched TCGA ADC short-read RNA-seq data for expression of *USFM*, but we could not find substantial read counts (<1 read per million) for samples with or without *U2AF1 S34F* mutations.

In contrast to gene sets identified from our differential isoform usage analysis, we found that differentially expressed isoforms belonged to genes regulated by NF-kB via TNF signaling, most of which are downregulated (corrected p-value <0.05; **Supplemental Table 6**). This observation is particularly important given recent reports implicating *U2AF1 S34F* in altering immune-related genes (Palangat *et al.*, 2019; Smith *et al.*, 2019). We further examined FLAIR isoforms derived from genes in the NF-kB pathway to determine if any *U2AF1 S34F*-associated splicing alterations could explain expression changes. However, when we overlapped results from our event-level splicing analyses along with our gene and isoform expression analyses we found no overlap between NF-kB affected isoforms and *U2AF1 S34F* altered splicing (**Figure 5B**). This result suggests that the expression of these isoforms may be modulated through a splicing-independent mechanism or the altered splicing event cannot be detected by our long-read data.

We expanded our alternative-splicing overlap analysis to ask which of the 198 isoforms with altered usage and expression coincided with other significantly altered features, such as alternative-splicing and gene expression. We found several clusters of features that partially explain the involvement of *U2AF1 S34F* mutation in isoform expression and usage dysregulation. For example, we found 27 isoforms with both significant isoform usage and cassette exon usage changes (**Figure 5C cluster C1**). It is possible that these isoforms with significant usage changes are defined by single exon skipping events and are likely directly induced by *U2AF1 S34F*. In contrast, we found several isoforms that did not overlap any other altered features (**Figure 5C clusters C2 & C7**). Similar to the isoforms from our NF-kb analysis, we suspect these isoform changes to be modulated through either a splicing-independent mechanism or splicing changes undetected by long-reads. We found a single gene, *UPP1*, that contained 4 overlapping features, which were changes in isoform expression, gene expression, cassette exon usage, and isoform usage. Altogether, we observe a consistent pattern of *UPP1* alterations associated with *U2AF1 S34F*, and also identify two populations of dysregulated isoforms that may be modulated through splicing-dependent and independent pathways.

**Figure 5.**
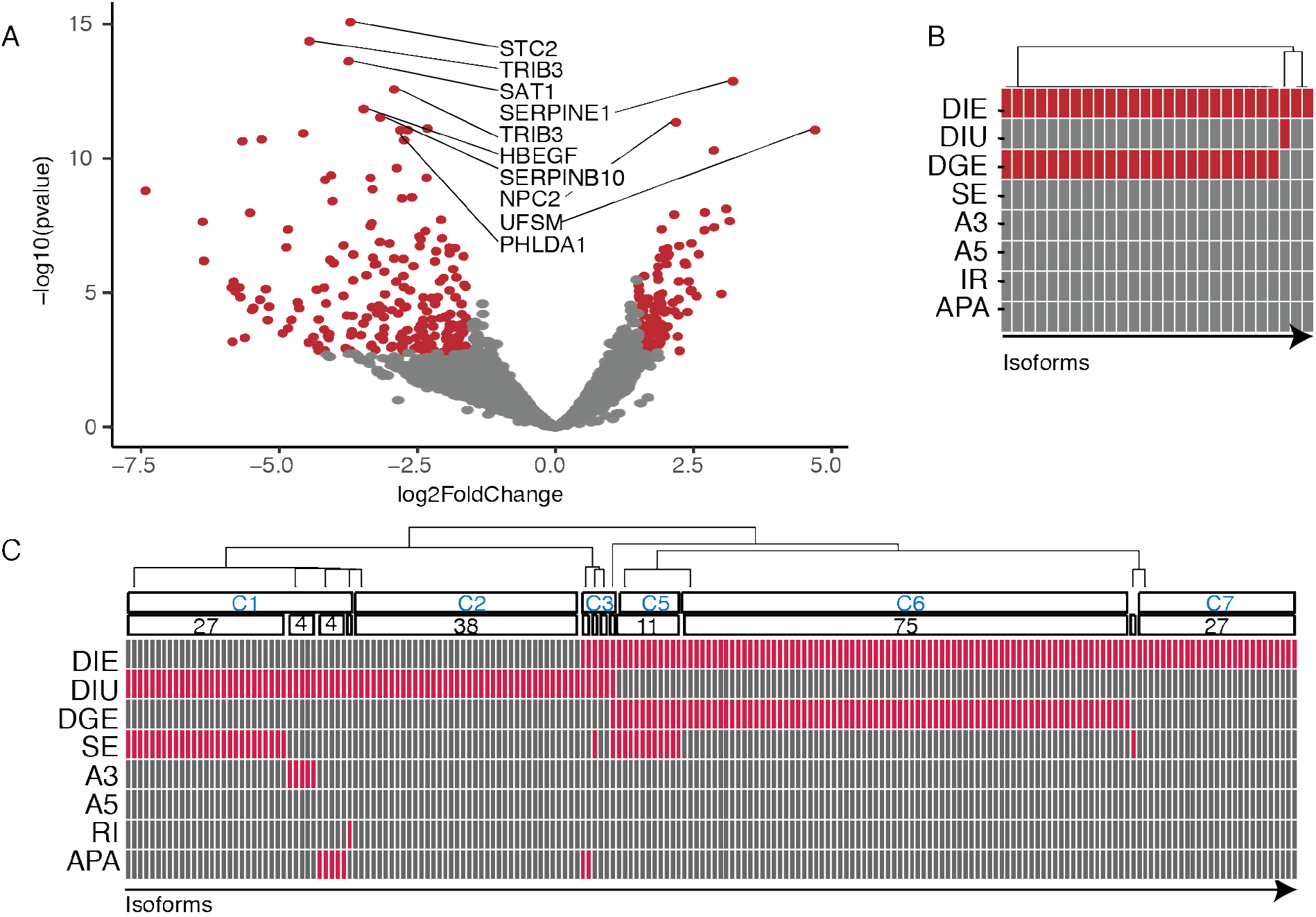
S34F-associated full-length isoform expression alterations. A) Volcano plot of differentially expressed isoforms. Red dots indicate expression changes with adjusted p-value<0.05 and magnitude change >1.5. Gene names are ordered by top 10 most significantly altered isoforms. B) Differential-event overlap of isoforms from genes involved NF-kb signaling pathway. Each box indicates an isoform, where red signifies if a particular isoform resulted as significantly altered for the corresponding analysis. DIE, differential isoform expression; DILI, differential isoform usage; DGE, differential gene expression; SE, skipped exon; A3, alternative 3’ splice site usage; A5, alternative 5’ splice site usage; IR, intron retention; APA, alternative polyadenylation site usage. C) Same as panel D, except including all isoforms with altered expression or usage.

### PTC-containing isoforms are downregulated by *U2AF1 S34F*

Our long-read approach enables a more confident open reading frame (ORF) prediction, which can be used to identify altered splicing events that are targeted by nonsense-mediated decay (NMD). NMD is a process that removes spliced mRNAs with premature termination codons (PTCs) that could give rise to gain-of-function or dominant-negative truncated protein products, if they were translated (Dreyfuss, Kim and Kataoka, 2002; Lewis, Green and Brenner, 2003; Sterne-Weiler *et al.*, 2013; Maslon *et al.*, 2014; Floor and Doudna, 2016; Aviner *et al.*, 2017). *U2AF1 S34F*-associated spliced products have been shown to be substrates of NMD (Yip *et al.*, 2017). Given the implications of U2AF1 S34F dysregulation and NMD, we asked transcriptome-wide what fraction of altered isoforms could be putative NMD targets. To do this, we classified FLAIR isoforms into two categories, either as putative protein-coding (PRO) isoforms or PTC-containing isoforms (**Methods**; **Figure 6A**). We postulated that the shallow sequencing depth of long-reads relative to short-reads would limit our ability in capturing PTC-containing isoforms if they are indeed subject to NMD. However, of our 63,289 FLAIR isoforms, we identified 8,037 PTC-containing isoforms (12% of all isoforms). We then asked what proportion of PTC-containing isoforms are dysregulated at the level of expression and isoform usage (**Figure 6B**). For differentially used isoforms, we found similar proportions of PTC-containing isoforms (Fisher’s exact two-sided test, p=0.5). In contrast, we found a significant difference in the proportion of PTC-containing isoforms between differentially expressed isoforms (Fisher’s exact two-sided test, p<0.01).

**Figure 6.**
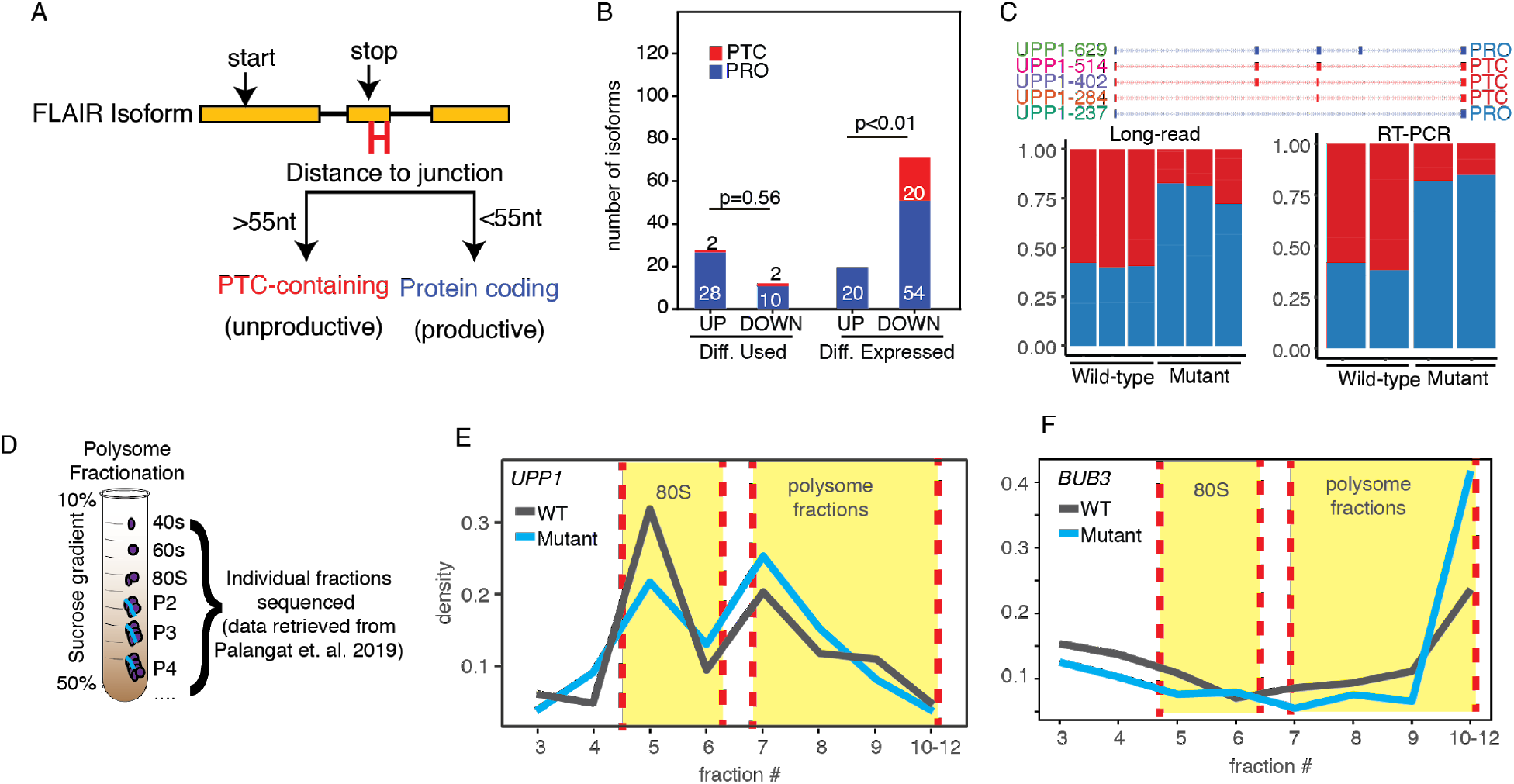
*U2AF1* S34F induces shifts in isoform productivity. A) Diagram of isoform productivity logic prediction. B) Comparison of up- and down-regulated S34F-associated isoform changes classified by productivity. UP, upregulated; DOWN, downregulated. Upregulated indicate isoforms with increased usage frequency or expression relative to wild-type. Downregulated indicate isoforms with decreased usage frequency or expression relative to wild-type. C) *UPP1* major isoforms classified by productivity. Bar plots show quantification of each productivity type for nanopore long-read data (left panel), and RT-PCR quantification (right panel). D) Polysome profiling analysis and sequencing scheme. E) *UPP1* expression density of normalized read counts across polysome fractions. Yellow highlights indicate 80s and polysome fractions. F) Same as panel E for *BUB3*.

We next sought to test, more generally, if *U2AF1 S34F* induces shifts in isoform productivity by conducting a gene-level analysis. To do this, we compared the proportion of PTC-containing versus productive isoform usage for each gene using the same methodology as our differential isoform usage analysis. Our results showed very few genes with strong shifts in productivity (**Supplemental Figure 4**; 10 total with corrected p-value <0.05 and ΔPTC isoform usage > 10%). However, we did identify a very strong shift in productivity in *UPP1 (*p-value<0.001 & Δproductivity > 20%), a gene we found to have strong changes in splicing, isoform usage, and expression. Interestingly, our differential gene expression analysis showed significant downregulation for *UPP1* (log2FoldChange 2.4 & p-value < 0.05), yet our productivity analysis showed a strong shift toward productive isoform usage (**Figure 6C**). Overall, our results suggest a bias toward downregulation of PTC-containing isoforms in *U2AF1 S34F* cells.

### *U2AF1 S34F* isoform dysregulation is associated with changes in translation

We predicted that if PTC-containing *UPP1* isoform are indeed subject to NMD, then the proportion of *UPP1* mRNAs able to undergo translation will be larger in mutant cells relative to wild-type, since there is a shift toward upregulation of productive *UPP1* isoforms. To test this, we used polysome profiling data from HBEC3kt cells with and without *U2AF1 S34F* causing mutations (**Methods; Figure 6D**). We found a significant change in the proportion of *UPP1* expression across different polysome fractions (chi-squared p-value < 0.01; **Figure 6E**). We observed a large drop (Δ10%) in the proportion of *UPP1* expression in polysome fractions 5 & 6 in the mutant relative to wild-type. These fractions correspond to the monosome, which is a fraction not associated with active translation, and is known to harbor non-coding mRNAs, such as NMD products (Floor and Doudna, 2016). The marked shift of *UPP1* expression in mutant samples from the monosome toward higher polysome (fractions >=7) is consistent with the hypothesis that *U2AF1 S34F*-associated *UPP1* alterations alter mRNA fate by shifting isoform production toward isoforms associated with enhanced translational activity.

We next tested if *U2AF1 S34F*-associated isoform changes in *BUB3* are consistent with differences in polysome profiles. In contrast to *UPP1*, we did not observe significant isoform productivity changes for *BUB3*. Instead, we observed significant changes in a terminal alternative 3’ splice site event that is linked to alternative polyadenylation site usage. Previous reports show that *BUB3* APA site 5 is associated with enhanced translational efficiency (Bava *et al.*, 2013). Our APA analysis showed mutant-specific isoform shifts toward isoforms with APA site 5, effectively increasing the proportion of translationally efficient *BUB3* isoforms. We tested for changes in *BUB3* polysome profiles using the same methodology used for *UPP1*. We find a strong shift in *BUB3* expression toward high polysome fractions (**Figure 6F**; chi-squared p-value <0.01). Notably, RNA-IP results from previous studies do not support large changes in cytosolic U2AF1 binding for *BUB3* or *UPP1*, which is a proposed mechanism of mutant *U2AF1* to modulate translational efficiency (Palangat *et al.*, 2019). Altogether, our data indicate a role for translational control through a splicing-dependent manner, and demonstrate distinct mechanisms of *U2AF1 S34F* for modulating translation control of genes through spliced isoform dysregulation.

We next determined if changes in translational control is a general feature for genes with strong changes in isoform expression and usage. Our results showed that 66% (42/63) of genes with *U2AF1 S34F*-associated isoform changes also had a significant change in polysome profile (**Methods**). This proportion was significantly higher (fishers two-sided test, p-value <0.01) than the 48% (1340/2753) of genes without S34F-associated isoform changes. Altogether, our results are consistent with previous work implicating *U2AF1 S34F* as a modulator of the translational landscape (Palangat *et al.*, 2019).

## Discussion

In this study, we assessed the impact of *U2AF1 S34F*-associated RNA processing alterations on individual mRNAs using an isogenic cell line harboring a *U2AF1 S34F* mutant allele. Although splicing alterations associated with *U2AF1* have been characterized with short-read sequencing, the full-length isoform context in which the altered events occur has not been described. We aimed to fill this gap in knowledge by using a long-read sequencing approach and supplemented our analysis with orthogonal short-read RNA-sequencing datasets from the same isogenic cell lines.

We demonstrate the robustness of long-read approaches by recapitulating splicing signatures associated with *U2AF1 S34F* mutations. Although our long-read transcriptome captures a comparable number of isoforms relative to short-read approaches, we still lack sequencing depth to capture the entire catalog of cassette exons associated with *U2AF1 S34F*, such as known cassette exons in *STRAP* or *ASUN* which were previously described to have *U2AF1 S34F*-associated splicing alterations (Fei *et al.*, 2016). Moreover, although we identified genes with significant changes in polyadenylation site selection, we were unable to recapitulate transcriptome-wide levels observed in previous studies (Park et. al. 2016). In line with these shortcomings, a saturation analysis of full-length isoform construction reveals isoform discovery limitations, possible due to relatively shallow sequencing depth (**Supplemental Figure 6**). However, long-read sequencing approaches offered by PacBio and Oxford nanopore are continually improving sequencing throughput and quality. Recent studies using newer Nanopore flow cell chemistry and higher-throughput platforms have demonstrated data yield orders of magnitude greater than this study (Tang *et al*., 2018). With greater data yield and improved transcriptome coverage, there is the potential to identify more *U2AF1 S34F* dysregulated isoforms with greater confidence.

We observe an interesting link between isoform dysregulation and translational control. Previous studies using RNA immunoprecipitation assays have shown that cytosolic mRNA binding of U2AF1 can modulate translational control (Palangat *et al.*, 2019). This splicing-independent mechanism of translational control is complementary to our findings here, in which isoforms arising from RNA processing alterations caused by *U2AF1 S34F* cause changes in translational control of the gene. Interestingly, our data supports two potential mechanisms. In the case of *BUB3*, *U2AF1 S34F* induces isoform switches toward isoforms with regulatory sequences that promote high translational efficiency. Alternatively, for *UPP1* we observe a substantial shift away from PTC-containing isoforms, which could serve as putative NMD targets. While further studies are necessary to directly test if these PTC-containing isoforms are regulated by NMD, we show that the expression of PTC-containing isoforms is strongly downregulated in the presence of *U2AF1 S34F*.

Our analyses contribute several findings implicating *UPP1* as severely dysregulated by *U2AF1 S34F*. So far, no reports have mentioned isoform-specific dysregulation associated with *UPP1*. *UPP1* has been observed to be upregulated in certain cancer types (Liu *et al.*, 1998). In our study using non-cancer derived cells, we find an opposite pattern, in which *UPP1* is significantly downregulated at the level of overall gene expression. The observed downregulation of *UPP1* is consistent with our finding of downregulation of isoforms involved in the TNF via NF-kB signaling pathway, which is a positive regulator of *UPP1* expression (Wan *et al.*, 2006). However, although we observe a strong downregulation at the level of total gene expression, our isoform usage and productivity analyses reveal a shift toward more productive isoforms. Nevertheless, further studies are required to determine what impacts *UPP1* isoform changes have on cellular function.

Overall, our data captured the context in which *U2AF1 S34F* RNA processing alterations occur at fulllength isoform resolution. We build upon previous short-read analyses by providing an extensive list of isoform-specific changes associated with *U2AF1 S34F*, along with the first estimates of isoform function. Our results demonstrate the importance of investigating the transcriptome of mutant splicing factors using long-read data that provides diverse perspectives on RNA processing and isoform function.

## Methods

### Data generation and processing

#### Preparing RNA for long-read sequencing

HBEC3kt cells with and without *U2AF1 S34F* were cultured as previously described (Ramirez *et al.*, 2004; Fei *et al.*, 2016). Total RNA was extracted from whole cell lysate using Zymo Direct-zol RNA kits. Purified RNA was prepared for long-read following previously established protocols (Picelli *et al.*, 2013; Byrne *et al.*, 2017; Tang *et al.*, 2018). Total RNA was reverse transcribed using the SmartSeq2 protocol, and amplified using 15 cycles of PCR. 1 ug of PCR amplified cDNA from each sample was subsequently used for Oxford Nanopore 1D library preparation (SQK-LSK108) on flow cell chemistry version 9.4. Basecalling was performed using Albabacore version 2.1.0 using options --flowcell FLO-MIN106 and --kit SQK-LSK108. Nanopore reads were prepared for genomic alignment by removing adapter sequenced using Porechop version 0.2.3 (Wick, 2017). After adapter removal, reads were aligned to GENCODE hg19 using minimap2 version 2.14-r894-dirty (Li, 2018) using the ‘-ax’ option.

#### Processing TCGA LUAD short-read data

Lung adenocarcinoma short-read data from The Cancer Genome Atlas (601 samples total) was downloaded from CGhub using gtdownload (Wilks *et al.*, 2014). TCGA donors with multiple RNA-seq bams were filtered by date to only include the most recent RNA-seq bam (495 samples). 495 TCGA bams were subsequently processed through JuncBase using default parameters with GENCODE hg19 comprehensive annotations and basic annotations as input to ‘getASEventReadCounts’ for options ‘ --txt_db 1’ and ‘txt_db2’, respectively (Brooks *et al.*, 2011). Differential splicing analyses were performed using Wilcoxon-rank sum between samples containing *U2AF1 S34F* splicing factor mutation (n=11) or no splicing factor mutation (n=451), which were defined by molecular profiling details outlined in Campbell et. al. (Campbell *et al.*, 2016).

#### Obtaining and processing HBEC3kt short-read data

Short-read HBEC3kt data was retrieved from NCBI short read archive (GSE80136). Reads were aligned to GENCODE hg19 using STAR version 2.5.3a (Dobin *et al.*, 2013) with parameters ‘--twopassMode Basic’. Aligned bams were subsequently individually used for transcriptome assembly using StringTie version 1.3.5 using GENCODE hg19 basic annotations (Pertea *et al.*, 2015). Individual GTF annotation files generated from StringTie were then merged using default parameters. For the differential splicing analysis of HBEC3kt short-read data, we used JuncBASE with the same methodology as described in TCGA LUAD short-read data methods section. HBEC3kt short-read data had two biological replicates per condition (wild-type and mutant); therefore, for statistical testing, we conducted pairwise fisher’s tests, then defined significant events as ones with a Benjamini-Hochberg corrected p-value > 0.05 within each condition and a corrected p-value <0.05 between samples across conditions. We then post-filtered significant events to remove redundant and overlapping events by running JuncBASE scripts ‘makeNonRedundantAS.py’ and ‘getSimpleAS.py’. To compare long and short-read Δpercent spliced-in values (PSI), we computed PSI changes for significant long-read cassette exon events by subtracting DRIMSeq-calculated proportion values for wild-type and mutant. We then filtered our short-read JuncBASE PSI table for significant long-read events, and computed the short-read change in PSI by subtracting the average PSI between wild-type and mutant.

### Long-read Analysis

#### Nanopore read correction, FLAIR-correct

Aligned Nanopore sequencing data were concatenated prior to running FLAIR v1.4 (Tang *et al.*, 2018) using samtools v 1.9 (Li *et al.*, 2009). Bam files were converted to bed using FLAIR-bam2bed12. Converted bed alignments were subsequently corrected using ‘FLAIR-correct’ with GENCODE hg19 basic annotations. Junctions identified by STAR alignment of HBEC3kt short-read data were also used as input into FLAIR-correct. Briefly, STAR junctions were kept if they contained at least 3 uniquely aligned in either both Mut1a and Mut1b samples or in both WT1 and WT2 samples. Junctions that did not follow GT-AG splicing motif were also removed.

#### FLAIR-collapse and diffExp

Differential analyses were performed by FLAIR-diffExp with default parameters. Genes and isoforms with less than 10 reads from either sample group were excluded from isoform expression and usage analyses.

#### Long-read alternative-splicing analysis, FLAIR-diffSplice

Differential alternative splicing for long-read data was conducted with FLAIR-diffSplice. FLAIR-diffSplice calls events for the following alternative-splicing types: cassette exon usage, alternative 3’ splice site, alternative 5’ splice site, intron retention, and alternative polyadenylation. Percent spliced-in values for each event were calculated by tallying the number of reads supporting isoforms that include an event, divided by the total number of reads that span the event. Inclusion and exclusion counts were then constructed into a table to process with DRIM-seq (Nowicka et. al. 2016) for differential splicing analysis.

#### Long-read alternative polyadenylation analysis

Poly(A) cleavage sites were defined by clustering FLAIR isoform transcript end sites using BedTools cluster, with a window distance of 5 (Quinlan and Hall, 2010; Quinlan, 2014). Poly(A) sites were then quantified by summing the total number of aligned read counts for each isoform that fell within each cluster. Clusters were assigned to genes, and counts for each cluster were then processed by DRIM-Seq. Genes with corrected p-value < 0.05 were considered to have significant changes in poly(A) site usage.

### Gene-set enrichment analysis

The Molecular Signatures Database (Liberzon *et al.*, 2011, 2015) was used to perform all gene set enrichment analysis using gene sets: GO gene sets, Hallmarks and Canonical pathways. Genes names included for isoform expression and isoform usage analyses were from isoforms with corrected p-value <0.05, and magnitude changes of ΔLog2FoldChange>1.5 and Δ10% isoform usage. Duplicate gene names from genes with multiple significantly altered isoforms were included only once.

### Polysome analysis

Polysome profiling data from HBEC3kt cells with and without *U2AF1 S34F* mutation were obtained from Palangat et. al. (2019; Supplemental Table S4). For each gene, normalized read counts across polysome fractions 3 through 10-12 were compared between mutant and wild type samples using Chi-squared test. Genes with less than 11 normalized read counts in any given fraction were not tested. Multiple testing correction was conducted using the python module statsmodels.stats.multitest.multipletests with default parameters. Significant changes in polysome profile were considered to have a corrected p-value of <0.05. We tested for general polysome profile alterations in *U2AF1 S34F*-associated genes by comparing the ratio of affected genes with and without significant changes in polysome profile versus unaffected genes. Affected genes were considered ones with either a significant isoform expression or usage change.

### Statistics and significance testing

Results from all differential analyses were called significant if their corrected p-value fell below p<0.05 and passed a magnitude filter. For differentially expressed isoforms, events over a log2 fold change of 1.5 were called significant. For differentially used isoforms and alternative splicing events, events with >=10% change in usage were called significant.

## Acknowledgments

We thank Dennis Fei and Harold Varmus for providing the HBEC3kt cells. We also thank the ENCODE Consortium and the ENCODE production lab of Bradley E. Bernstein and the Kellis Computational Biology group for production of the histone regulatory data used via the UCSC Genome browser tracks for this study. This work was supported by the Pew Charitable Trusts and the University of California Tobacco-Related Disease Research Program T29KT040I (A.N.B.). A.D.T. was funded through NIH grant T32HG008345 and R01HG010053. C.M.S. was supported by training grants NIH T32GM008646, R25GM058903, and the Ford Foundation predoctoral fellowship.

## Author Contribution

C.M.S. and A.N.B. designed the study. C.M.S. and E.H.R designed experiments. C.M.S performed experiments. C.M.S., A.D.T., M.G.M., A.N.B. wrote code and analyzed data. C.M.S., E.H.R., A.D.T., A.N.B. interpreted the data. C.M.S. and A.N.B. wrote the manuscript with input from all other co-authors.

## Code Availability

All FLAIR related scripts and modules used in this study can be found at https://github.com/BrooksLabUCSC/FLAIR. FLAIR commands and other code are available as jupyter notebooks upon request.

## Data Availability

Long-read nanopore sequencing data from HBEC3kt wild type and *U2AF1 S34F* cells are available in the NCBI GEO database (GSE140734 accession number).

