## Supplemental Information for "Nanopore sequencing reveals *U2AF1 S34F*-associated full-length isoforms"

for

**Supplemental Figures**

Supplemental Figure 1

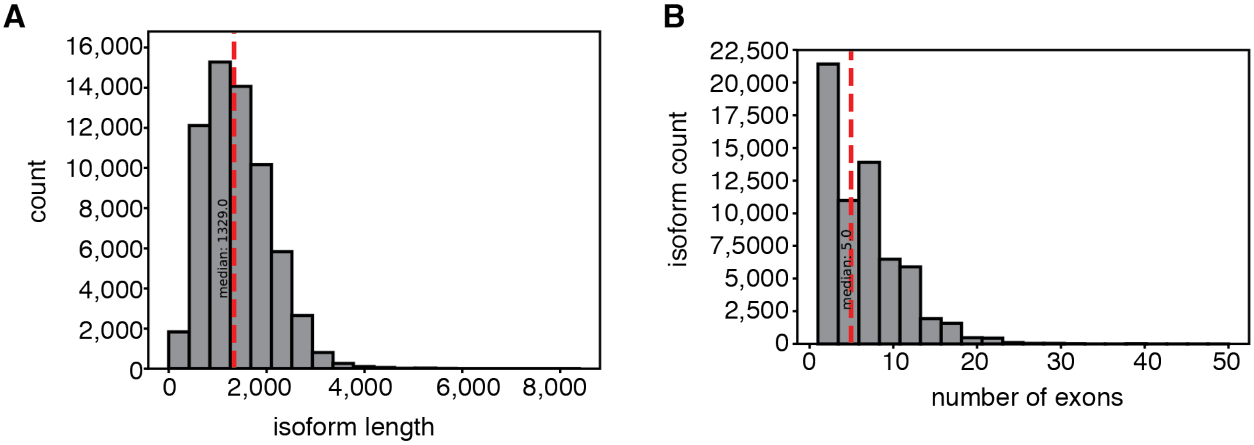


**Supplemental Figure 1 | FLAIR isoform lengths and exon numbers.** A) Isoform lengths for FLAIR collapsed and filtered isoforms. Red dashed line indicated isoform length median B) Distribution of exon numbers per FLAIR collapsed and filtered isoforms.

Supplemental Figure 2

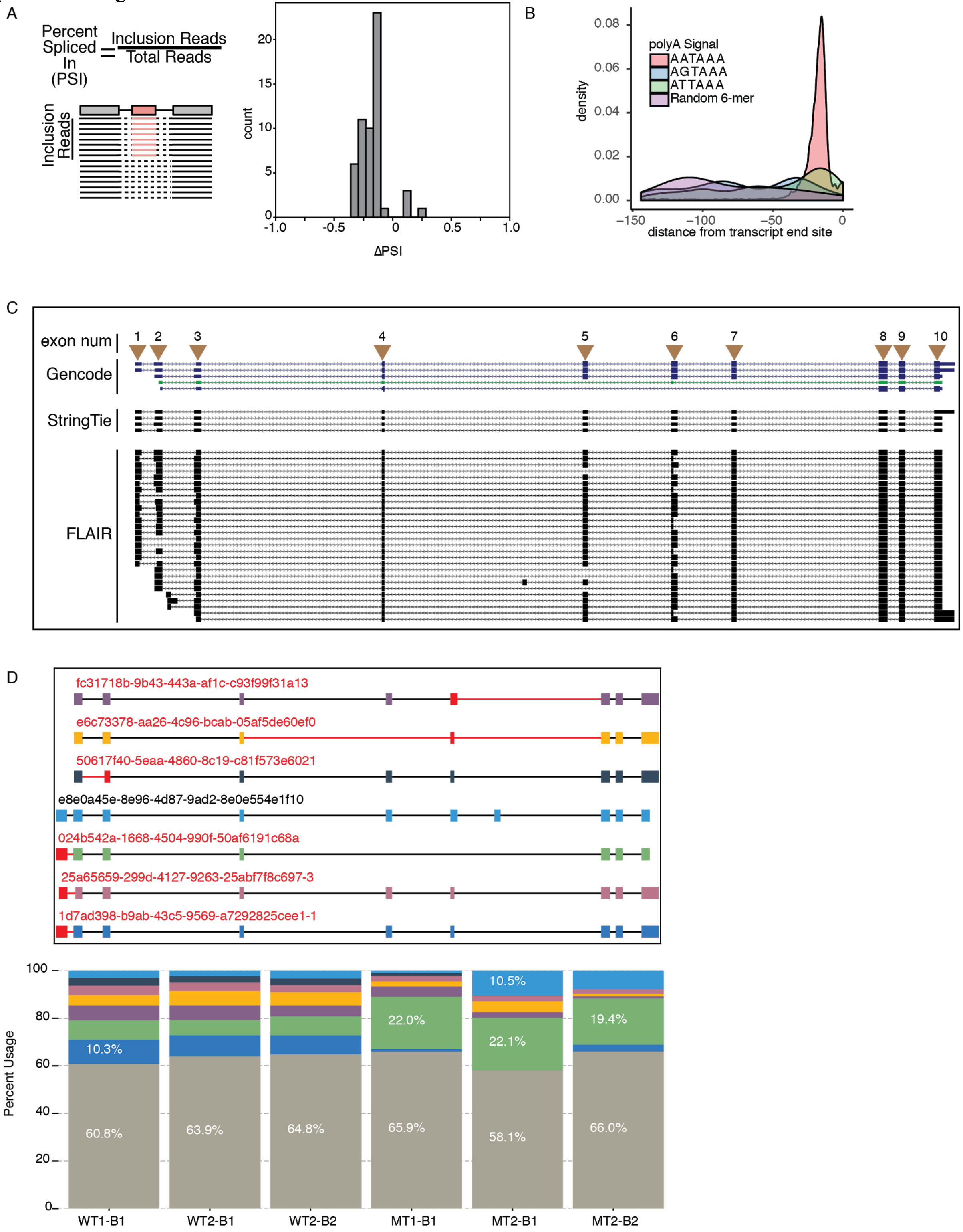


**Supplemental Figure 2 | Event-level long-read analyses.** A) Percent spliced-in computation (left panel) and histogram showing overall change in percent spliced in values between mutant and wild-type. B) Poly(A) cleavage site k-mer analysis on long-read FLAIR isoforms. C) UCSC genome browser shot of *UPP1* showing a diagram StringTie and FLAIR isoforms containing exon 7. D) *UPP1* top 7 expressed isoforms (top panel) and long-read quantification (bottom panel). Red isoform names indicate isoforms not assembled by short-reads, and red intron/exon indicate novel connectivity. Gray bars in stacked bar plot indicate isoforms expressed less than 5%.

Supplemental Figure 3

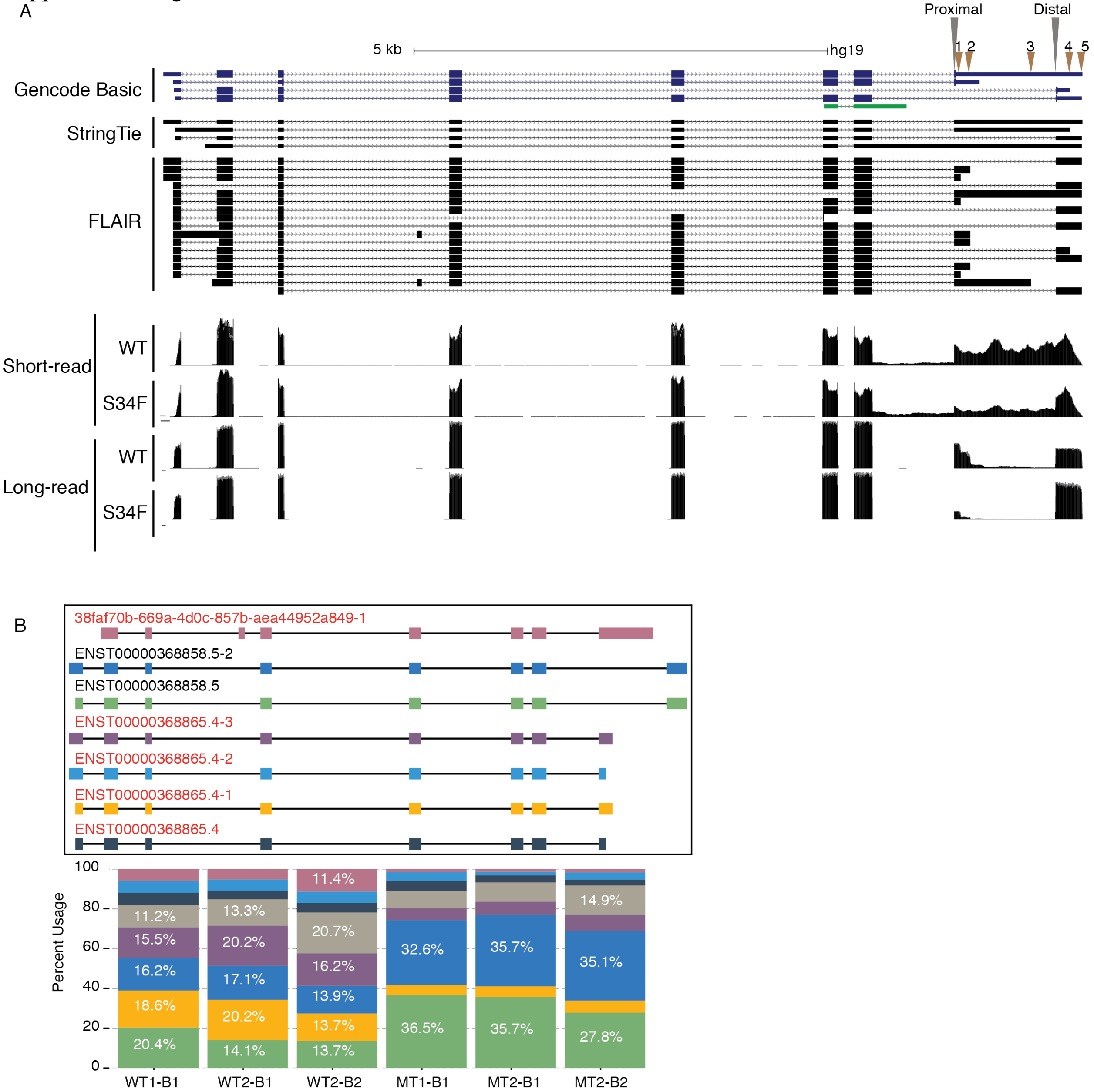


**Supplemental Figure 3 | *BUB3* isoform switching is associated with *U2AF1 S34F.*** A) *BUB3* isoforms and coverage data. Top three panels show isoforms from GENCODE v19, StringTie assembly, and FLAIR, respectively. Bottom four panels show sequencing data coverage plots from short-read and long-read data, from wild-type (WT) and *U2AF1 S34F* (S34F) samples. Yellow highlights indicate alternative-splicing events that define *BUB3* isoforms. Brown arrows indicate alternative polyadenylation sites. B) Top 7 most expressed BUB3 isoforms (left panel), and quantification for each isoform (top panel). Red isoform names correspond to isoforms not present in short-read assembly. Color in bottom panel correspond to isoform quantification.  Gray color in stacked bar plot represent quantification with less than 5% usage.

Supplemental Figure 4

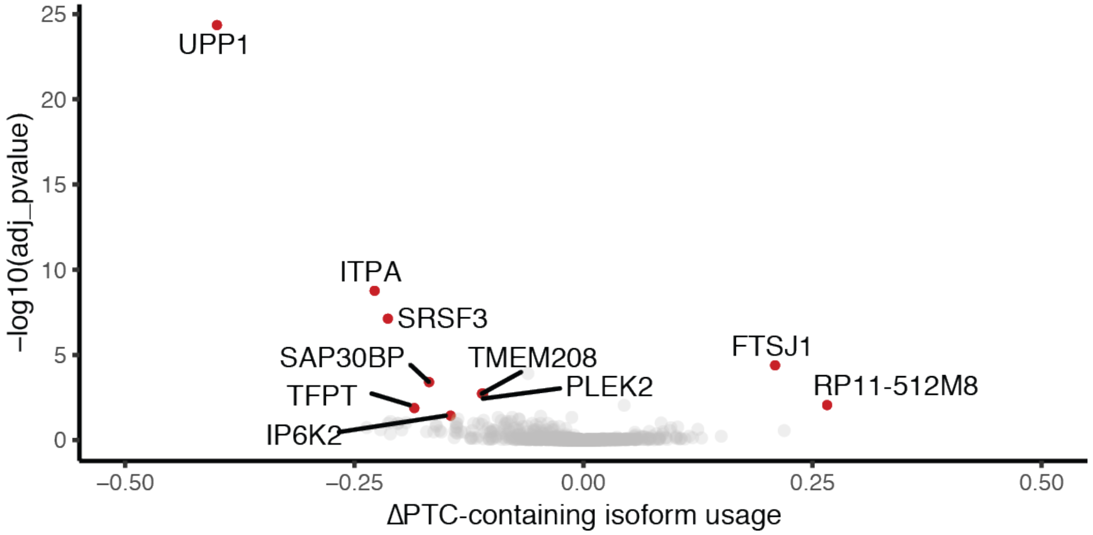


**Supplemental Figure 4 |** Volcano plot of differential isoform productivity analysis. Red points indicate genes with productivity shifts where corrected p value < 0.05 and change in productivity is greater than 10%. Gray points indicate genes with corrected p value >= 0.05.

Supplemental Figure 5

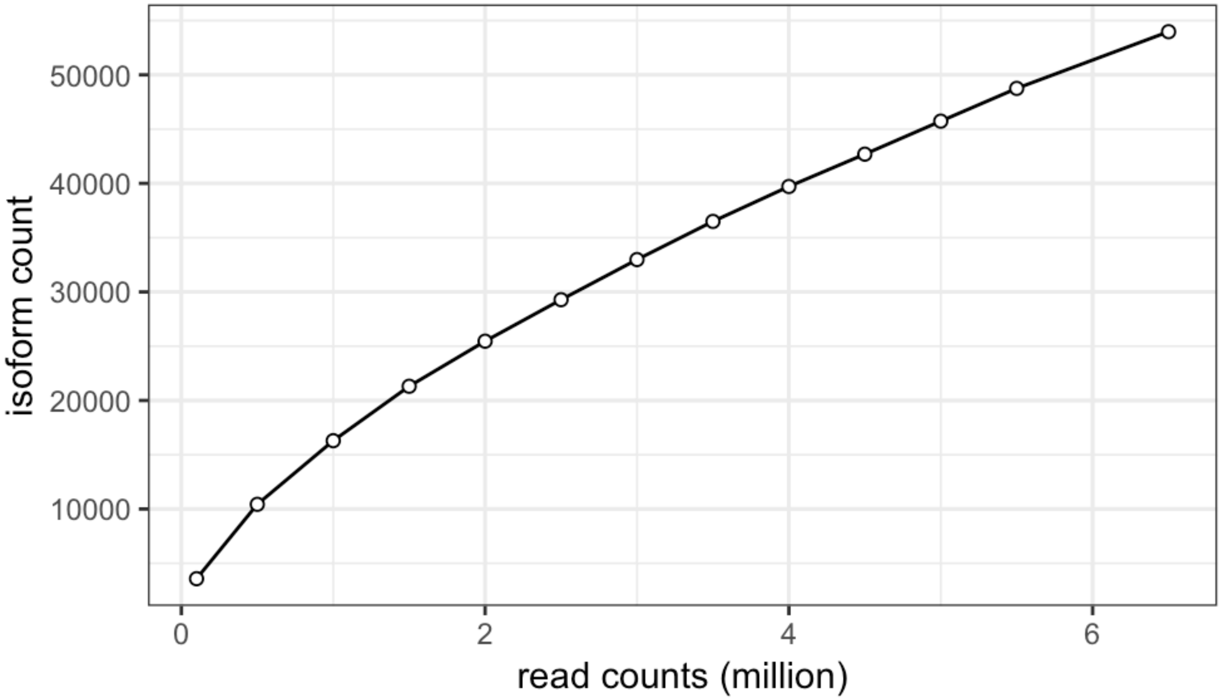


**Supplemental Figure 5 | Isoform identification saturation curve.** Reads from all 6 sequencing libraries were concatenated and subsampled at various depths: 0.5-6.5 million reads at intervals of 0.5m

Supplemental Tables

**Supplemental Table 1 - Nanopore read length and GC statistics.** Tab-delimited matrix containing statistics for each sequencing run for WT1, WT2, MT1, MT2  samples (biological replicate 1 and 2 - B1 & B2). Columns describe the following: numReads - total number of reads, numBases - total number of bases called for each sequencing run, meanLen - read length statistical average , medLen - read length statistical median, minLen - minimum read length, maxLen - longest read length, meanGC - GC content statistical average, medGC - GC content statistical median.

**Supplemental Table 2 - JuncBASE table of TCGA-associated U2AF1 splicing events.** Tab-delimited matrix containing alternative-splicing events and quantifications from short-read TCGA lung adenocarcinoma RNA-seq data identified by juncBASE. Each entry represents an individual splicing event. Each column represents distinct characteristics for each event: novel_event - describes whether the inclusion coordinate overlaps GENCODE v19 annotated intron boundaries, as_type - describes the alternative-splicing pattern type, hugo - denotes the hugo symbol gene name for each event, chromosome & strand - describe the chromosome and reference event strand, exclusion_coords & inclusion_exon_coord - define the genomic ranges which exclude and include each event, ΔPSI - is the difference in PSI between U2AF1 S34F mutant and non-mutant samples, p-value & p.adj - are the raw and benjamini hochberg adjusted p-values from wilcoxon rank sum test.

**Supplemental Table 3 - JuncBASE table of short-read HBEC3kt analysis.** Tab delimited matrix containing alternative-splicing events and quantifications from short-read HBEC3kt samples from Fei et. al. (2016). Samples are denoted by SRA sample numbers. Columns describe event-type, inclusion and exclusion coordinates (similar to Supplemental Table 2). Values in columns headed by samples SRR numbers denote percent spliced in (PSI) values range from 0 to 100. Events for which a sample did not have at least 25 supporting reads are denoted as “NA”

**Supplemental Table 4 - List of significant FLAIR-diffSplice of splicing events.** Tab delimited file of alternative splicing events identified in long-read data. Alternative-splicing type and coordinate of event are described in the first column: es - cassette exon, a3 - alternative acceptor, a5 - alternative donor, ir - retained intron. Remaining columns describe the magnitude change in percent spliced in (PSI) as determined by DRIM-seq and significance of change.

**Supplemental Table 5 - List of differentially used FLAIR isoforms and enriched gene sets.**
Table 1 corresponds to differentially used isoforms as determined by DESeq2. Log2FoldChanges represent shinkage computed changes as computed by the LFCShrinkage function. Isoform names not conforming to ENSEMBL transcript names denote novel isoforms, and gene names not conforming to ENSEMBL gene names correspond to novel gene loci. Table 2 corresponds to results from gene set enrichment using gene names from used isoforms.

**Supplemental Table 6 - List of differentially expressed genes and FLAIR isoforms.** Same as Supplemental Table 5, but for differentially expressed genes and isoforms. Table 1 corresponds to differentially expressed isoforms as determined by DESeq2. Table 2 corresponds to results from gene set enrichment using gene names from differentially expressed isoforms. Table 3 corresponds to differentially expressed genes.

**Supplemental Files**

**Supplemental File 1 - GTF of FLAIR isoforms.** General transfer formatted (GTF) list of mRNA isoforms identified by FLAIR.

**Supplemental File 2 - GTF of StringTie isoforms.** Same as Supplemental File 1 but for isoforms assembled by StringTie.

**Supplemental File 3 - FLAIR isoform count table.** Tab delimited file containing raw expression from FLAIR-quantify. Columns correspond to isoform name and sample ids.
